# Wnt/β-catenin signaling and p68 conjointly regulate CHIP in colorectal cancer

**DOI:** 10.1101/2021.06.20.449107

**Authors:** Satadeepa Kal, Shrabastee Chakraborty, Subhajit Karmakar, Mrinal K Ghosh

## Abstract

The differential expression pattern of Carboxy terminus of Hsc70 Interacting Protein (CHIP, alias STIP1 Homology and U–box Containing Protein 1 or *STUB1*) in cancers is associated with ubiquitination mediated degradation of its client proteins. Emerging evidences suggest its abundant expression of CHIP in colorectal cancer compared to normal tissues, but the mechanistic detail of this augmented expression pattern is unclear. The signature driver of canonical Wnt pathway, β-catenin, and its co-activator RNA helicase p68, are also overexpressed in colorectal cancer. In this study, we describe a novel mechanism of Wnt/β-catenin and p68 mediated transcriptional activation of CHIP gene leading to enhanced proliferation of colorectal cancer cells. Wnt3A treatment and pharmacological activation of canonical Wnt signaling pathway resulted in increased nuclear translocation of β-catenin and elevated expression of CHIP. Likewise, overexpression and knockdown of β-catenin and p68 upregulated and downregulated CHIP expression, respectively, at both mRNA and protein levels. After cloning CHIP promoter, the increased and decreased promoter activities of CHIP induced by overexpression and knockdown of either β-catenin or p68 further confirmed transcriptional regulation of CHIP gene by Wnt/β-catenin signaling cascade. p68 along with β-catenin were found to occupy Transcription Factor 4 (TCF4) binding sites on endogenous CHIP promoter and regulate its transcription. Finally, enhanced cellular propagation and migration of colorectal cancer cells induced by ‘Wnt/β-catenin-p68–CHIP’ axis established the significance of this pathway in oncogenesis. To the best of our knowledge, this is the first report elucidating the mechanistic details of transcriptional regulation of CHIP (*STUB1*) gene expression.

## INTRODUCTION

In recent years, colorectal cancer has emerged as one of the leading cancers in terms of morbidity and fatality. It is the third most important cause of cancer worldwide and second in causing mortality ^1^. Thus, insightful studies of the mechanism of the enhancement or diminution of this cancer have gained immense importance. Colorectal cancer has often been hallmarked by the aberrant increase or decrease of different proteins wired to help in the progression of the disease. One such factor is β-catenin, a crucial transcription regulator involved in the canonical Wnt/β-catenin signaling pathway that aids in cellular proliferation and poor prognosis by promoting the transcription of several target genes ^2^. Mutations in β-catenin regulatory pathway lead to its reduced destruction, increased accumulation, and subsequent migration to the nucleus to bind to transcription factor TCF4 at TCF4 binding elements (TBE) ^3^. This binding promotes oncogenesis by increased transcription of proto-oncogenes Cyclin D1 ^4^, c-myc ^5^, AKT ^6^. Similarly, p68, a typical RNA helicase of the DDX (DEAD Box) family, is often observed to be upregulated in multiple cancers, functioning as co-activator of several transcription factors like β-catenin, Androgen receptor (AR), STAT3, p53, NF-κB, and Estrogen Receptor alpha (ERα) ^7^. In colorectal cancer, it shows high expression and a prominent association with β-catenin in the nucleus, enhancing transcription of β-catenin driven genes ^8^.

Not only the transcription factors but Ubiquitin ligases also dictate the stability of proteins during oncogenesis ^9,10^ and hence are important determinants for the fate of the disease. One such ubiquitin ligase is CHIP, encoded by the *STUB1* (STIP1 Homology And U–box Containing Protein 1) gene, which is known to be differentially expressed in multiple cancers ^11^. Recent reports suggest that CHIP is overexpressed in colorectal cancer samples compared to normal colon ^12,13^. Established as a co-chaperone, CHIP performs a significant role in maintenance of protein homeostasis by aiding in degradation of a plethora of cellular proteins in various diseases ^14^. In cancer, it regulates the expression of proteins like p53 ^15^, PTEN ^16^, p21 ^17^, ERα ^18^, c-myc ^19^, Arc ^20^, which are classified as either oncogenes or tumor suppressors. However, very less is known about the regulation of CHIP apart from a few post transcriptional, post translational, and protein-protein interaction based mechanisms. Post-transcriptional modification of CHIP by miR-764-5p inhibits its translation and affects osteoblast differentiation ^21^, whereas its post translational modification by Cdk5 promotes neuronal death ^22(p5)^. Aurora Kinase A mediated phosphorylation of CHIP protein aids in enhanced degradation of AR in prostate cancer ^23^. Interaction with Ca^2+^/S100 proteins also inhibits E3 ligase activity of CHIP ^24^. While the above findings state the regulation of E3 ligase activity of CHIP protein, no report explicates its transcriptional regulation in any context.

Consequently, a thorough elucidation of the transcriptional regulation of CHIP gene could explain the context dependent expression of the protein. Since colorectal cancer is associated with an aberrant escalation in the canonical Wnt/β-catenin signaling and p68, as mentioned earlier, along with an increased amount of CHIP protein ^12^, we intended to perform a detailed mechanistic study on whether the CHIP gene is transcriptionally regulated by the Wnt/β-catenin signaling cascade with possible involvement of p68. This study provides us with the initial insights into the transcriptional regulation of CHIP (*STUB1*) gene expression in the context of colorectal cancer. Here, we have cloned the promoter of CHIP for the first time and uncovered a novel mechanism by which β–catenin and p68 transcriptionally regulate its expression, facilitating cancer progression through cell proliferation and migration.

## MATERIALS AND METHODS

### Bioinformatics analyses

*STUB1* gene expression was analyzed in multiple cancers including colorectal cancer using UALCAN database. A Heat map was generated using Human Protein Atlas. The expression of *STUB1* in different datasets of colorectal cancer was further checked in Oncomine data. The specimens were compared as cancer vs normal patients. p value <0.05 was considered significant.

### Cell culture, Transfection and Treatments

HCT 116 was maintained in McCoy’s 5A media. HT29, SW480 and SW620 and HEK 293 were maintained in Dulbecco’s Modified Eagle’s Medium (DMEM) supplemented with 10% Fetal Bovine Serum (FBS) and antibiotic Penicillin, Streptomycin, and Gentamycin at a recommended dose as previously described [25]. DNA constructs were transfected using Lipofectamine 2000 (Invitrogen). siRNA mediated silencing of p68, TCF4 and CHIP were executed using Lipofectamine RNAimax (Invitrogen) for 48 hours. The siRNA transfections were performed for 48 hours. In case of drug treatment following transfection of DNA, transfections were done at least 24 hours earlier to drug treatment. LiCl (Calbiochem), the agonist of β-catenin pathway, and FH535 (Sigma Aldrich), the antagonist, were administered at 30mM for 4 hours and 15uM for 24 hours, respectively upon standardization of the doses. Recombinant Human Wnt 3a (BioVision) was administered at 150 ng/ml for 24 hours. LiCl and Wnt3a were added to previously serum-starved cells.

### Cloning and plasmid constructs

Promoter sequence of *STUB1* gene was obtained from Eukaryotic Promoter Database (EPD). A region spanning 2163 bps upstream from the transcription start site was PCR amplified using genomic DNA isolated from HEK293 cells with Platinum SuperFi II DNA polymerase (Thermo Fisher Scientific). The PCR product was cloned into pGL3 basic vector following conventional protocols. The primers used for cloning are mentioned in Appendix A. The putative binding sites of the transcription factors on the promoter were studied from ALGGEN-PROMO (http://alggen.lsi.upc.es/). β-catenin and p68 genes were sub-cloned from pBI and pSG5 plasmids into pGZ21dx vector for expression of GFP-β-catenin and GFP-p68, GFP-tagged fusion proteins. CHIP was cloned into pIRES-hrGFP1a vector. The DNA constructs were further ascertained by restriction digestion and sequencing. The list of the primers used is given in Appendix A.

### Small interfering RNA (siRNA) and short hairpin RNA (shRNA) mediated knockdown

shRNA against β-catenin was purchased from Addgene (pLKO.1 puro shβ-catenin; Addgene cat no.18803). Scramble siRNA, CHIP siRNA, TCF4 siRNA and p68 siRNA were bought from Santa Cruz Biotechnology (Santa Cruz Biotechnology, Inc., Dallas, Texas, USA) and added at a final concentration of 60nM.

### Site directed mutagenesis

Deletion mutants of two TCF4 binding sites (TBE1 and TBE2) on the *STUB1* promoter were generated as previously performed [25] using the Quickchange XL Site directed mutagenesis kit (Agilent technologies, Santa Clara, CA, USA) as per the manufacturer’s protocol. These were sequenced and the primer sequences are given in Appendix A.

### Immunoblotting

Whole cell lysates, nuclear and cytoplasmic extracts were prepared and immunoblotting was executed as described in previous studies ^25,26^. About 40ug of lysates were subjected to SDS-Polyacrylamide Gel Electrophoresis, followed by transfer to PVDF membrane, blocking (1 hour) with 5% BSA, incubation with primary antibodies (overnight at 4°C) and HRP-conjugated secondary antibodies (2 hours at room temperature), before developing the blots with Millipore Luminata Classico. The primary antibodies used were β-catenin, GFP, GAPDH, CHIP, Lamin B (Santa Cruz Biotechnology); TCF4, Cyclin D1(Cell Signaling Technologies), p68 (Abcam), Actin (Sigma Aldrich). HRP tagged anti-goat, anti-mouse, and anti-rabbit secondary antibodies were purchased from Sigma Aldrich. The densitometric scanning and quantification of blots were done using GelQuant.Net software.

### Preparation of RNA and quantitative Real time PCR

TRIZOL Reagent (Invitrogen) was used to isolate total RNA as per the manufacturer’s protocol. 1ug of RNA was converted to cDNA using High-capacity cDNA transcription kit (Thermo Fisher scientific), and consequently used for quantitative Real time PCR analyses using SyBr Green master mix (Biorad) in ViiA 7 Real-time PCR Instrument (Applied Biosystems). The Standard deviation calculation and quantification were done from three independent experiments. 18s rRNA was used as the internal control (for normalization). The primer sequences used are given in Appendix A.

### Luciferase assay

The cells were transfected with PGL3-*STUB1* promoter construct or its mutants together with Renilla luciferase construct (pRL-TK). Promoter activity was analyzed according to the experiments mentioned in the figures. The luciferase assay was performed after 48 hours using a Dual luciferase assay kit (Promega, Madison, WI, USA). The results were quantified from three independent biological repeats. The values were quantified luminometrically by the Varioskan Flash Multimode Reader (Thermo fisher scientific, Waltham, MA, USA) as described previously ^27^.

### Chromatin immune precipitation (ChIP) assay

Preparation of the chromatin fragments and subsequent immune precipitation were accomplished as described previously ^25^. Briefly, 1% paraformaldehyde was used to crosslink HCT 116 cells for 10 minutes at room temperature. The cross-linking was quenched by 0.125M glycine in rocking condition. The cells were washed with PBS thrice and pelleted. After resuspension in ChIP lysis buffer, they were sonicated to a fragment size of ~500bp. The fragmented chromatin was centrifuged, and the lysate was precleared with pre-blocked (in 5% BSA) Sepharose A bead. 10% of the lysate was kept aside as input after pre-clearing. The samples were incubated with 3ug of primary antibodies against p68 (Abcam), β-catenin, TCF4, Polymerase II or normal Rabbit IgG (Santa Cruz Biotechnology, Inc., Dallas, Texas, USA) at 4°C, overnight and pulled down further using blocked Sepharose A beads. The precipitated chromatin was washed with wash buffers and eluted from the beads. Thereafter de-crosslinking was done. The DNA obtained after de-crosslinking and purification was PCR amplified by TopTaq DNA polymerase master mix (Qiagen), using the appropriate set of primers and run on a 2% agarose gel for visualization. The sequences of the primers are as given in Appendix A.

### Immunofluorescence microscopy

HCT 116 cells were cultured on coverslips placed on 35 mm culture plates until ~70% confluency. After performing treatments along with respective controls, the cells were fixed in 4% paraformaldehyde, permeabilized with 0.5% Triton-X-100, blocked with 3% BSA in PBS, washed with PBS, and subjected to immunostaining using primary antibodies against CHIP, β-catenin (Santa Cruz Biotechnology, Santa Cruz, SA, USA) and fluorescence conjugated secondary antibodies Alexafluor488 (AF488) and Alexafluor594 (AF594). The nuclei were stained with 4,6-Diamidino-2-phenylindole (DAPI). The images were captured at 60X by FluoView FV10i confocal laser scanning microscope (Olympus Life Science).

### Cell viability assay

Cell viability assay using MTT [3-(4,5-dimethylthiazol-2-yl)-2,5-diphenyl tetrazolium bromide] was done after cells were treated or transfected as per experimental design and cell confluency reached 70-80% as described previously ^28^. The absorbance of the colored product was measured at 550nm by VarioSkan Flash multimode reader (Thermo Fischer Scientific, Waltham, MA).

### Wound healing (scratch) assay

HCT 116 cells grown to 50-60% confluency in 35 mm plates, transverse scratches were made using 10 ul pipette tip at the beginning of transfections or treatments. Images were captured in Life Technologies EVOS L microscope after 24 hours of treatment and after 48 hours of siRNA-mediated transfection, as it takes longer to exhibit siRNA effects ^26^.

### Colony formation assay

Treatments and transfections were performed on HCT 116 cells seeded at a density of 1×10^3^, as per experimental design and with proper controls. After the respective time periods for treatments and transfections, cells were allowed to grow for 15 days in complete medium. The assay was performed as described previously ^29^.

### Cell cycle analysis

After transfection or treatment as per experiment, HCT 116 cells were harvested with Trypsin, washed in PBS, fixed with 70% ethanol with mild vortexing and kept in ice for 30 minutes. After further PBS washing, cells were resuspended in Propidium Iodide/RNase staining buffer (BD Biosciences) in dark. The cells were passed through a sieve to remove clumps and collected in FACS tubes before performing experiments in BD LSR-Fortessa using FACS Diva software (Version 8.0.2) of BD Biosciences.

### Statistical analyses

The statistical significance was drawn by conducting Two Samples Student’s t-test [27]. The significance values were represented as *P≤ 0.05, **P≤0.005, ***P≤0.0005, ****P≤0.00005 and the non-significant values were represented as NS. The statistical analyses were performed using GRAPHPAD PRISM8.

## RESULTS

### Bioinformatic analyses of CHIP (STUB1) expression

To generate an overall expression profile for *STUB1* in cancer, analysis of *STUB1* mRNA expression data in 24 different cancer types available in the The Cancer Genome Atlas (TCGA) database was performed using UALCAN ^30^, a web-based tool that facilitates in-depth analysis of TCGA database. The findings were plotted as log_2_(TPM+1) vs TCGA samples (Figure 1A) after data normalization in normal and tumor samples. In many of the cancer types, *STUB1* expression was found to be upregulated in tumor samples as compared to the respective normal samples. From these cancer types, Colorectal Adenocarcinoma (COAD) was chosen for further analysis. From COAD dataset, *STUB1* transcript levels were checked in primary tumor vs normal samples as well as in different stages of cancer progression. An increase in *STUB1* expression level was observed in primary tumor samples compared to the normal samples, and also with increasing grades of colorectal cancer progression (Figure 1B). To corroborate this finding with other repositories, a heatmap was generated from the pathological data globally available in The Human Protein Atlas (HPA) ^31^ also representing a high degree of *STUB1* expression in colorectal cancer (Figure 1C).

**FIGURE 1:**
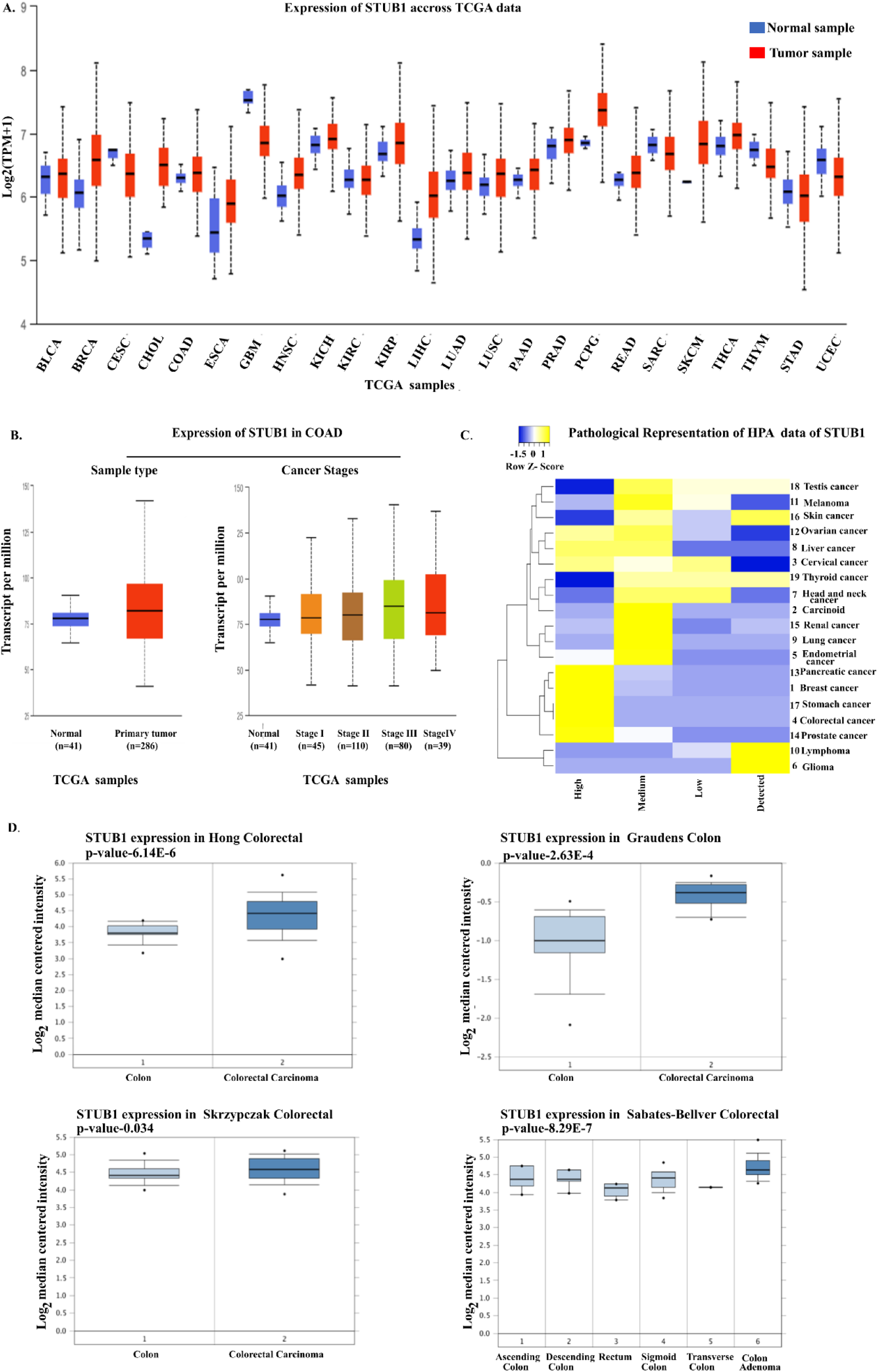
Gene expression profile of CHIP (*STUB1*). *A,* Bioinformatics based Pan-Cancer analysis of *STUB1* expression across the cancers using the UALCAN (http://ualcan.path.uab.edu/analysis.html) platform. BLCA, bladder urothelial carcinoma; BRCA, breast carcinoma; CESC, cervical squamous cell carcinoma and endocervical adenocarcinoma; CHOL, cholangiocarcinoma; COAD, colon adenocarcinoma; ESCA, esophageal carcinoma; GBM, glioblastoma multiforme; HNSC, head and neck squamous cell carcinoma; KICH, Kidney chromophobe; KIRC, kidney renal clear cell carcinoma; KIRP, kidney renal papillary cell carcinoma; LIHC, liver hepatocellular carcinoma; LUAD, lung adenocarcinoma; LUSC, lung squamous cell carcinoma; PAAD, pancreatic adenocarcinoma; PRAD, prostate adenocarcinoma; PCPG, pheochromocytoma and paraganglioma; READ, rectum adenocarcinoma; SARC, sarcoma; SKCM, skin cutaneous melanoma; THCA, thyroid carcinoma; THYM, thymoma; STAD, stomach adenocarcinoma; UCEC, uterine corpus endometrial carcinoma. *B,* The *STUB1* gene expression level in tumor tissues samples compared with that in normal tissue samples (left panel) and a cancer stage specific (I-IV) study of *STUB1* expression level of tumor samples in comparison with normal tissue samples in UALCAN analysis (right panel). *C,* Heatmap showing the pathological expression profiles of CHIP (*STUB1)* protein across different cancer type, data available at HPA (https://www.proteinatlas.org). The yellow-colored clusters represent high expression in Z-score, and the blue-colored clusters represent low expression in Z-scores. *D,* Analysis of *STUB1* gene expression in adjacent normal colon vs colorectal cancer samples in Oncomine (https://www.oncomine.org/resource/login.html). The box plot comparing region specific *STUB1* expression in normal colon (left plot) and colorectal cancer samples (right plot) was derived from Oncomine database. The analysis shown were for Hong Colorectal, Skrzypczak Colorectal, Graudens Colon and in Sabates-Bellver Colorectal datasets.

For further validation of this observation, Oncomine ^32^ analysis of *STUB1* gene expression was performed in selected colon cancer datasets, by comparing clinical specimens of cancer with normal patient datasets in each case (Figure 1D). Results with p value <0.05 were considered significant. This analysis confirmed that *STUB1* was significantly overexpressed in different datasets of colorectal cancer. Thus, multiple web-based servers indicated a higher expression of *STUB1* in colorectal cancer, which paved the way for discovering the underlying mechanism of its expression in this cancer.

### Canonical Wnt signaling enhances CHIP expression in colorectal cancer

An elevated level of CHIP protein in colorectal cancer has been reported previously ^12^. Likewise, an aberrant overexpression of Wnt/β-catenin pathway proteins has also been widely observed in colorectal cancer ^33^. p68, a known transcriptional co-activator of β-catenin ^34^, is also known to be overexpressed in colorectal cancer ^35^. To uphold the preceding findings, and investigate any possible relationship among these proteins, the endogenous protein levels of CHIP, β-catenin, and p68 were checked in multiple colorectal cancer cell lines (Figure 2A). High expression of these proteins led to the use of HCT 116 and HT29 cell lines for further experiments. Wnt3A, a known ligand and inducer of canonical Wnt signaling pathway, was used for enhanced nuclear translocation of β-catenin and stimulation of this pathway ^36^. On treating HCT 116 cells with purified Wnt3a protein, we observed an increase in both protein and mRNA levels of CHIP along with an increased β-catenin protein level. Cyclin D1 was used as a positive control ^37^ for the pathway (Figure 2B). LiCl, a pharmacological inhibitor of GSK3β and known activator of β-catenin pathway ^38^, was used to enhance Wnt signaling in HCT 116 and HT29 cells. LiCl treatment at increasing concentrations of 10mM, 20mM, 30mM depicted a gradual increase in CHIP protein level along with β-catenin level in HCT 116 cells (Figure S1A). Next, the maximum concentration of 30mM of LiCl was used to treat HCT 116 and HT29 cells and an augmented protein level of β-catenin, CHIP, and Cyclin D1 was observed. (Figure 2C). In agreement with the above, we also see enhanced level of CHIP protein in both the cytosolic and nuclear components that correlates with increased nuclear migration of β-catenin upon LiCl treatment (Figure 2D). This is further endorsed by confocal microscopy study which illustrated an enhanced level of CHIP protein along with β-catenin nuclear translocation (Figure 2E). This suggests that enhanced β-catenin level due to activation of canonical Wnt signaling pathway is associated with an increase in CHIP level in colorectal cancer cells. To check whether the reverse holds true, FH535, an established inhibitor of the Wnt/β-catenin pathway ^39^ was used to diminish Wnt signaling pathway. With increasing concentrations of 5uM, 10uM, 15uM of FH535, there was a gradual decrease in CHIP protein level, along with β-catenin (Figure S1B). The highest concentration at which the decrease of CHIP protein was maximum, 15uM, was then used to treat both HCT 116 and HT29 cells. Protein levels of CHIP, Cyclin D1, and β-catenin decreased in both the cell lines upon inhibiting the pathway by this treatment (Figure 2F). HCT 116 cells were further treated with either LiCl or FH535 or both in conjunction and it was observed that CHIP protein levels positively correlated with enhancing the β-catenin pathway and decreased upon inhibiting the pathway. Interestingly, when the activator LiCl was administered on cells already treated with the inhibitor FH535, CHIP levels were seen to be rescued in the inhibitor-treated cells (Figure 2G). All these findings together suggest that the canonical Wnt signaling pathway involving β-catenin dictates the expression of CHIP in colorectal cancer cells.

**FIGURE 2.**
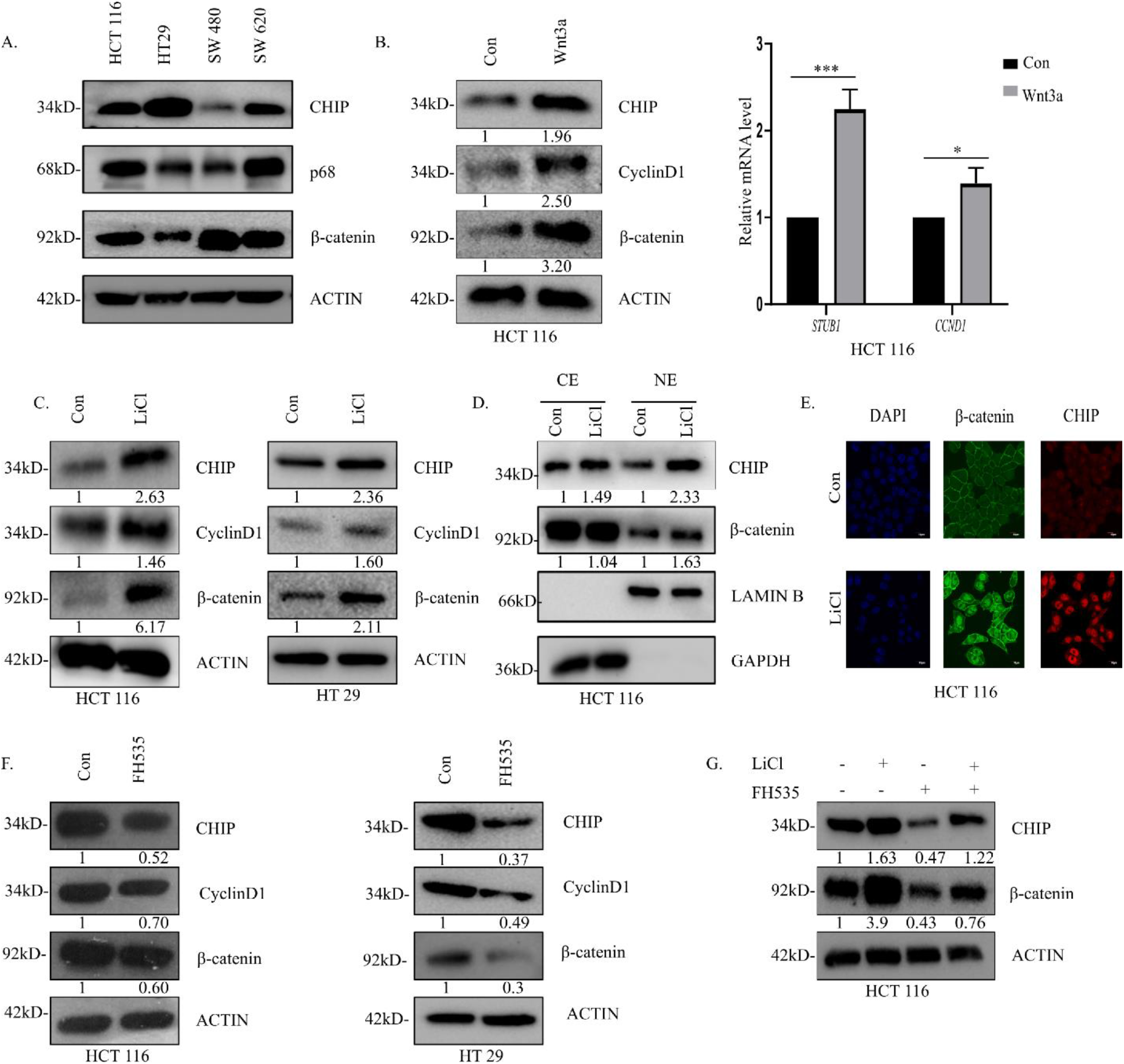
Canonical Wnt signaling enhances CHIP expression in colorectal cancer. *A*, Whole cell lysates prepared from HCT 116, HT29, SW480, and SW620 cells were immunoblotted with antibodies against CHIP, p68, and β-catenin. *B,* HCT 116 cells were treated with 150 ng/ml of purified Wnt3a for 24 hours. Whole cell lysates were probed with antibodies against CHIP, Cyclin D1, and β-catenin (left). cDNA prepared from the Wnt3a-treated HCT 116 cells was subjected to qPCR and transcript levels of *STUB1* (CHIP) and *CCND1* (Cyclin D1) (right) were checked. *C,* HCT 116 (left) and HT29 (right) cells were treated with 30mM LiCl for 4 hours. The whole cell lysates were probed for CHIP, Cyclin D1, and β-catenin. *D,* Control or LiCl-treated HCT 116 cells were subjected to subcellular fractionations. Both cytoplasmic extract and nuclear extract were immunoblotted with CHIP and β-catenin antibodies. *E,* Control or LiCl-treated HCT 116 cells were stained with primary antibodies against β-catenin and CHIP, subsequently secondary antibodies conjugated to AF488 (β-catenin, green) or AF594 (CHIP, red), were added and observed under fluorescent microscope. DAPI staining was done to visualize nuclei of the cells. The images were acquired at 60X magnification in an Olympus microscope. Scale bar-10um. *F,* HCT 116 (left) and HT29 (right) cells were treated with 15uM FH535 for 24 hours. The whole cell lysates were probed for CHIP, Cyclin D1, and β-catenin. *G,* HCT 116 cells were treated with either 30mM LiCl, or 15uM FH535, or both. Whole cell lysates were immunoblotted for CHIP and β-catenin. The experiments were carried out at least three times. *Con:* Control, *CE:* Cytoplasmic Extract, *NE:* Nuclear Extract. Densitometric values for immunoblots were calculated according to the loading control. Actin for whole cell lysates, Lamin B and GAPDH in case of cytoplasmic and nuclear extract were kept as loading controls. Error bars in all the designated subfigures represent mean (+) s.d. from independent biological repeats. The p values were calculated using Student’s t-test and p≤0.00005 is represented as **** for significant values.

### β-catenin regulates CHIP gene expression

β-catenin acts as a transcriptional coactivator by binding to the TCF/LEF complex enhances the transcription of its target genes ^40^. In colorectal cancer, the abnormal increase of its target genes that lead to poor prognosis of cancer makes it a prominent target for therapeutics ^41^. To understand whether β-catenin regulates CHIP expression in colorectal cancer, β-catenin was overexpressed in HCT 116 and HT29 cell lines. A heightened expression of CHIP due to overexpression of β-catenin was observed along with an increase in Cyclin D1 level, which acted as a positive control for β-catenin target genes (Figure 3A). A dose dependent study of β-catenin overexpression showed an increase in CHIP protein level with increasing dosage of β-catenin inside the cell (Figure 3B). To investigate further whether the increase in CHIP protein is a consequence of its increase at the mRNA level by β-catenin mediated transcription, CHIP mRNA levels were quantified under normal and β-catenin overexpressed conditions in HCT 116 and HT29 cells. In both the cell lines, an increase in mRNA level of CHIP was observed (Figure 3C). Cyclin D1 mRNA levels served as a positive control. On knocking down endogenous β-catenin levels, a decline in the protein level of CHIP and Cyclin D1 was observed in HCT 116 and HT29 cell lines (Figure 3D). Similar results were observed at the mRNA level, where upon downregulation of endogenous β-catenin levels, CHIP and Cyclin D1 mRNA levels declined as well (Figure 3E). β-catenin is known to interact with TCF4 to mediate transcription of target genes ^3^. Accordingly, on depletion of TCF4 by siRNA, a sharp decline in CHIP protein level was detected (Figure 3F). These observations collectively highlight that β-catenin positively regulates the transcription of CHIP gene which is also reflected at the protein level in colorectal cancer cell lines.

**FIGURE 3.**
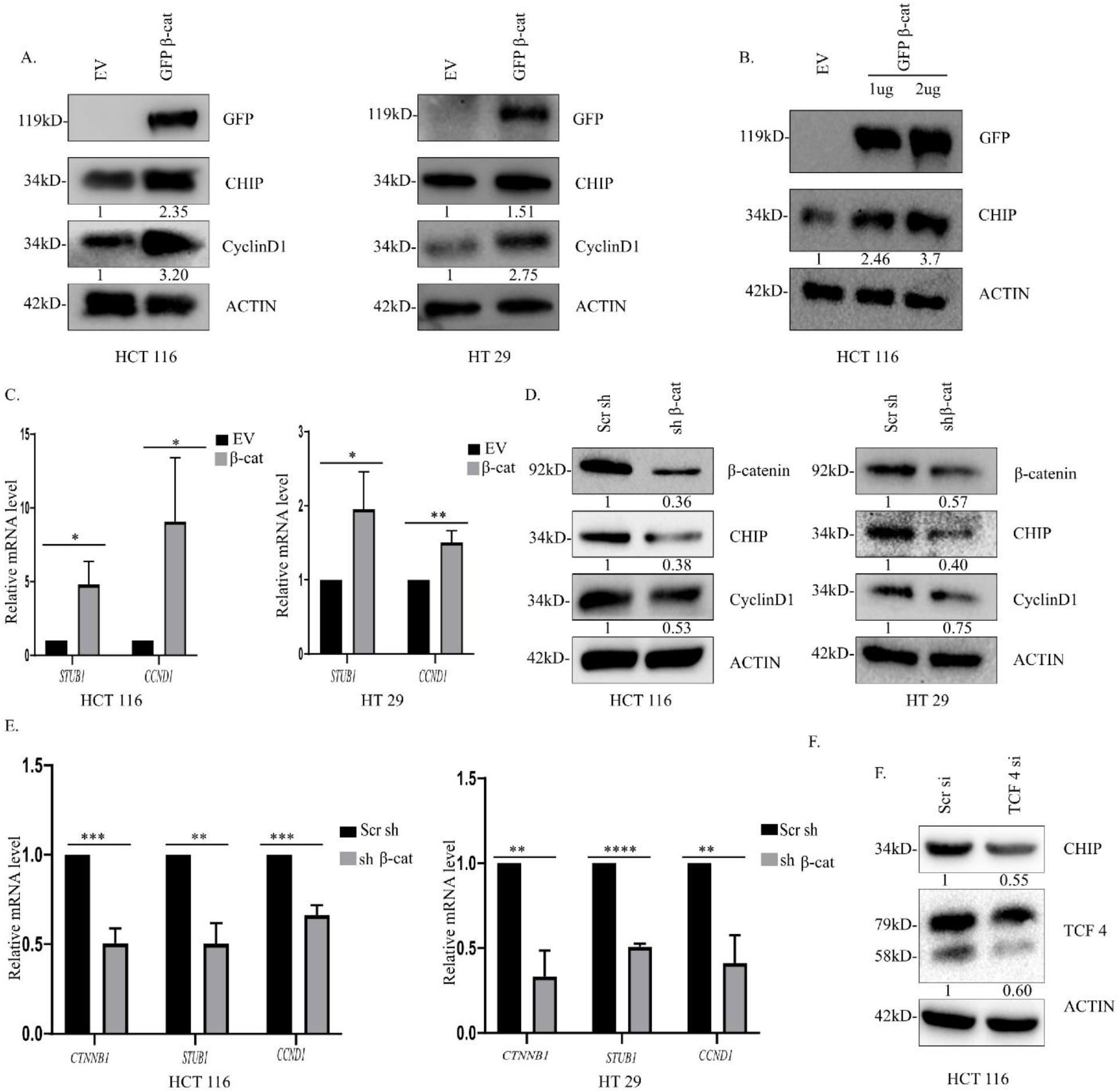
β-catenin regulates CHIP gene expression. *A,* HCT 116 (left) and HT29 (right) cells were transfected with 2ug of either empty vector (EV) or GFP-β-catenin. Whole cell lysates were probed for GFP (exogenous β-catenin), CHIP, and Cyclin D1. *B,* HCT 116 cells were transfected with either empty vector, or 1ug and 2ug of GFP-β-catenin. Whole cell lysates were immunoblotted with antibodies against GFP (exogenous β-catenin) and CHIP. *C*, cDNA prepared from the EV and β-catenin-transfected HCT 116 (left) and HT29 (right) cells was used for qPCR to check the transcript levels of *STUB1* (CHIP) and *CCND1* (Cyclin D1) for positive control. *D,* HCT 116 (left) and HT29 (right) cells were transfected with 2ug each of scrambled shRNA or β-catenin shRNA. Whole cell lysates were immunoblotted with antibodies against β-catenin, CHIP, and Cyclin D1. *E,* qPCR was done with cDNA prepared from the β-catenin shRNA-transfected HCT 116 (left) and HT29 (right) to check the transcript levels of *CTNNB1* (β-catenin), *STUB1* (CHIP) and *CCND1* (Cyclin D1). *F,* HCT 116 cells were transfected with 2ug of either scrambled siRNA or TCF4 siRNA. Whole cell lysates were prepared and probed for CHIP and TCF4. The experiments were conducted at least three times. *GFP-β-cat:* GFP-β-catenin, *Scr sh:* Scrambled shRNA, *Scr si:* Scrambled siRNA, *shβ-cat:* β-catenin shRNA. Densitometric values for immunoblots were calculated with respect to the loading control Actin. Error bars in all the designated subfigures represent mean (+) s.d. from independent biological repeats. Indicated p values were calculated using Student’s t-test and p≤0.00005 is represented as ****.

### RNA helicase p68, a transcriptional co-activator of β-catenin/TCF4 complex, additionally controls CHIP gene expression

p68, a co-activator of β-catenin ^34^ aids in transcription of β-catenin target genes and is known to be upregulated in colorectal cancer ^42^. To explore any possible role of p68 in expression of CHIP in colorectal cancer, p68 was overexpressed in HCT 116 and HT29 cells. Immunoblotting results revealed an increased level of CHIP protein in p68 overexpressed lanes (Figure 4A, 4B). Cyclin D1 acted as the positive control ^43^. To check whether the regulation is at the transcriptional level, cDNA prepared from empty vector transfected control and p68 overexpressed cells were subjected to quantitative real time PCR. An increase of CHIP mRNA level was also observed under p68 overexpressed condition, along with increased expression of Cyclin D1 mRNA (Figure 4A, 4B). A dose dependent study of p68 overexpression in HCT 116 resulted in an increase in CHIP protein level with increasing doses (Figure 4C). Further, on knocking down endogenous p68 level by siRNA targeted against p68, a concomitant decrease in CHIP protein level was noted, along with a decrease in Cyclin D1 in HCT 116 and HT29 cell lines (Figure 4D, 4E). A similar decrease in CHIP mRNA level was also observed upon p68 knockdown, along with a decrease in p68 and Cyclin D1 mRNA level (Figure 4D). Thus, p68 further controls CHIP gene expression in colorectal cancer cells.

**FIGURE 4:**
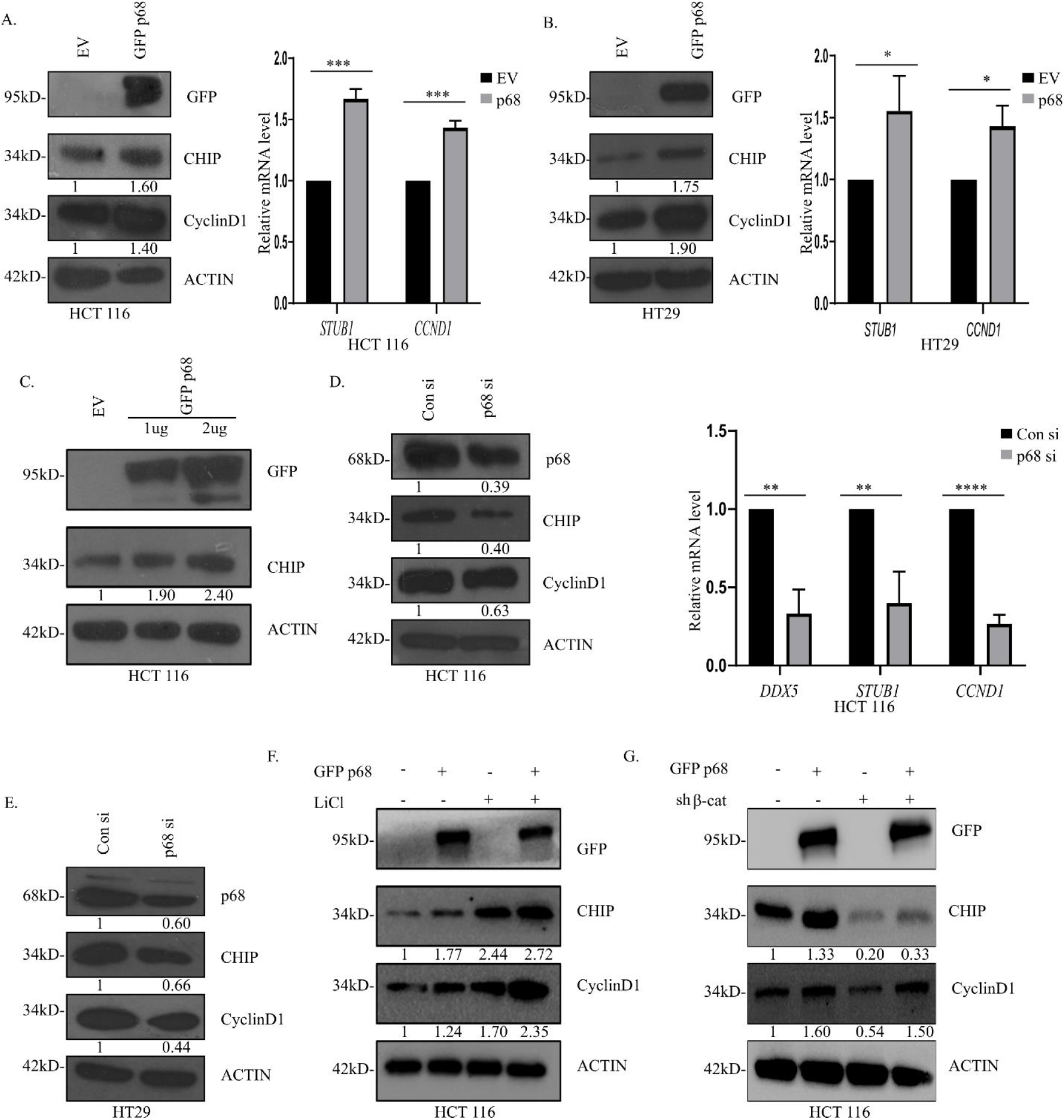
RNA helicase p68, a transcriptional co-activator of β-catenin/TCF4 complex additionally controls CHIP gene expression. *A,* HCT 116 cells were transfected using 2ug of either EV or GFP-p68. Whole cell lysates were probed for GFP (exogenous p68), CHIP, and Cyclin D1 (left). cDNA prepared from control and p68-transfected HCT 116 cells were analyzed for the change in transcript levels of *STUB1* (CHIP) and *CCND1* (Cyclin D1) (right) by qPCR. *B,* HT29 cells were transfected with 2ug of either EV or GFP-p68. Whole cell lysates were probed for GFP (exogenous p68), CHIP, and Cyclin D1 (left). cDNA prepared from the p68-transfected HT29 cells was subjected to qPCR to check the transcript levels of *STUB1* (CHIP) and *CCND1* (Cyclin D1) (right). *C,* HCT 116 cells were transfected with either EV, or 1ug or 2ug of GFP-p68. Whole cell lysates were immunoblotted with antibodies against GFP (exogenous p68) and CHIP. *D,* HCT 116 cells were transfected with 2ug of either Scrambled siRNA or p68 siRNA. Whole cell lysates were immunoblotted with antibodies against p68, CHIP and Cyclin D1 (left). cDNA prepared from the control and p68 siRNA transfected HCT 116 cells was subjected to qPCR to check the transcript levels of *DDX5 (*p68), *STUB1* (CHIP) and *CCND1* (Cyclin D1) (right). *E,* HT29 cells were transfected with 2ug of either Scrambled siRNA or p68 siRNA. Prepared whole cell lysates were immunoblotted with antibodies against p68, CHIP, and Cyclin D1. *F,* HCT 116 cells were subjected to either transfection with 2ug GFP-p68, or treatment with 30mM LiCl (4 hours), or both. Whole cell lysates were probed for GFP (exogenous p68), CHIP and Cyclin D1. *G,* HCT 116 cells were subjected to either transfection with 2ug GFP-p68, or transfection with 2ug β-catenin shRNA, or co-transfection with both. Whole cell lysates were probed for GFP (exogenous p68), CHIP and Cyclin D1. The experiments were carried out at least three times. *EV:* Empty Vector, *Scr si:* Scrambled siRNA. Densitometric values for immunoblots were calculated with respect to the loading control Actin. Error bars in all the designated subfigures represent mean (+) s.d. from three independent biological repeats. The p values were calculated using Student’s t-test and p≤0.00005 is represented as ****.

To delve deeper into the collective function of β-catenin and p68 in CHIP gene regulation, LiCl treatment and p68 overexpression were performed on HCT 116 cells, either individually or in conjunction. It was observed that CHIP protein level increased upon either p68 overexpression or administration of LiCl and was further augmented when both were administered together (Figure 4F). In the next experiment, β-catenin knockdown and p68 overexpression were performed either individually or in conjunction. While knockdown of β-catenin led to a decrease in CHIP protein and overexpression of p68 led to its increase, an intermediary level of CHIP protein was observed when both were administered in HCT 116 cells (Figure 4G). These observations cumulatively state that the CHIP gene expression is regulated by β-catenin, which is also aided by the transcriptional co-activator, p68.

### Transcriptional activation of CHIP (STUB1) promoter is under the control of both β-catenin and p68

To investigate whether the recruitment of β-catenin and p68 occurs at the promoter of *STUB1*/CHIP gene, further studies were conducted. The CHIP promoter sequence was retrieved from Eukaryotic Promoter Database. When analyzed for the putative sites of the transcription factors that might bind to the CHIP promoter, two TCF binding elements (TBEs) spanning regions −2033 to −2026 and −1985 to −1977 were found on the CHIP promoter. The sites were numbered according to their position upstream of the transcription start site. A schematic of the TBEs on the CHIP promoter is represented (Figure 5A). Since β-catenin and its co-activator p68 act *via* TBEs, a chromatin immunoprecipitation (ChIP) experiment was conducted in endogenous condition in HCT 116 cells to check for promoter occupancy using anti-TCF4, anti-p68, and anti-β-catenin antibodies. The region spanning the TBE sites on CHIP promoter was amplified using the primers (mentioned in Appendix A) spanning both the sites and the amplified product was named as T1. The data revealed binding of β-catenin, p68, and TCF4 to the CHIP promoter. The TBE regions of Cyclin D1 promoter were also amplified, serving as a positive control. GAPDH was taken as a negative control (Figure 5B). Next, ChIP experiment was conducted in LiCl treated and control cells. The promoter occupancy was observed to be much more in case of LiCl treatment thus proving that β-catenin occupied the CHIP promoter more when the pathway gets activated (Figure 5C). This observation justifies the enhanced expression of CHIP upon activation of Wnt/β-catenin signaling pathway (Figure 2C). Once the promoter occupancy was established, the role of β-catenin and p68 in altering the promoter activity needed to be checked. A 2163 bps fragment of the CHIP promoter was cloned into PGL3 basic vector, the schematic representing the insert sequence with highlighted forward and reverse primers (Figure 5D). To the best of our knowledge, this is a maiden report of the cloning of CHIP promoter. A positive clone retrieved after cloning was subjected to restriction enzyme digestion with the respective sites. The intact plasmid of size 6981 bps, when digested, yielded a 4818 bps PGL3 vector and a 2163 bps promoter insert, all of which were run on an agarose gel and sequenced to confirm a proper clone (Figure 5E). After successful cloning of the wild-type promoter, deletion mutants were constructed with individual deletions of the two TBE sites. The promoter construct with a deletion in the region −2033 to −2026 was termed as ΔTBE1 mutant and that with deletion in the region −1985 to −1977 was termed as ΔTBE2 mutant, as represented in the schematic (Figure 5F). The promoter activity measured by dual luciferase assay revealed an increase upon β-catenin overexpression and a decrease when β-catenin is downregulated (Figure 5G). Luciferase assay after β-catenin overexpression with the wild type and deletion mutant promoters proved that on deletion of the TBEs, there was a significant reduction in the activity of the CHIP promoter (Figure 5H). When similar experiments were conducted with p68 overexpression and knockdown, increased p68 levels elevated the promoter activity of CHIP and a decrease in p68 endogenous levels led to the opposite result (Figure 5I). Thus, these observations in conjunction state that upon activation of Wnt/β-catenin signaling pathway, β-catenin and its co-activator p68 mediate transcriptional activation of the CHIP promoter and regulate CHIP gene expression in colorectal cancer.

**FIGURE 5.**
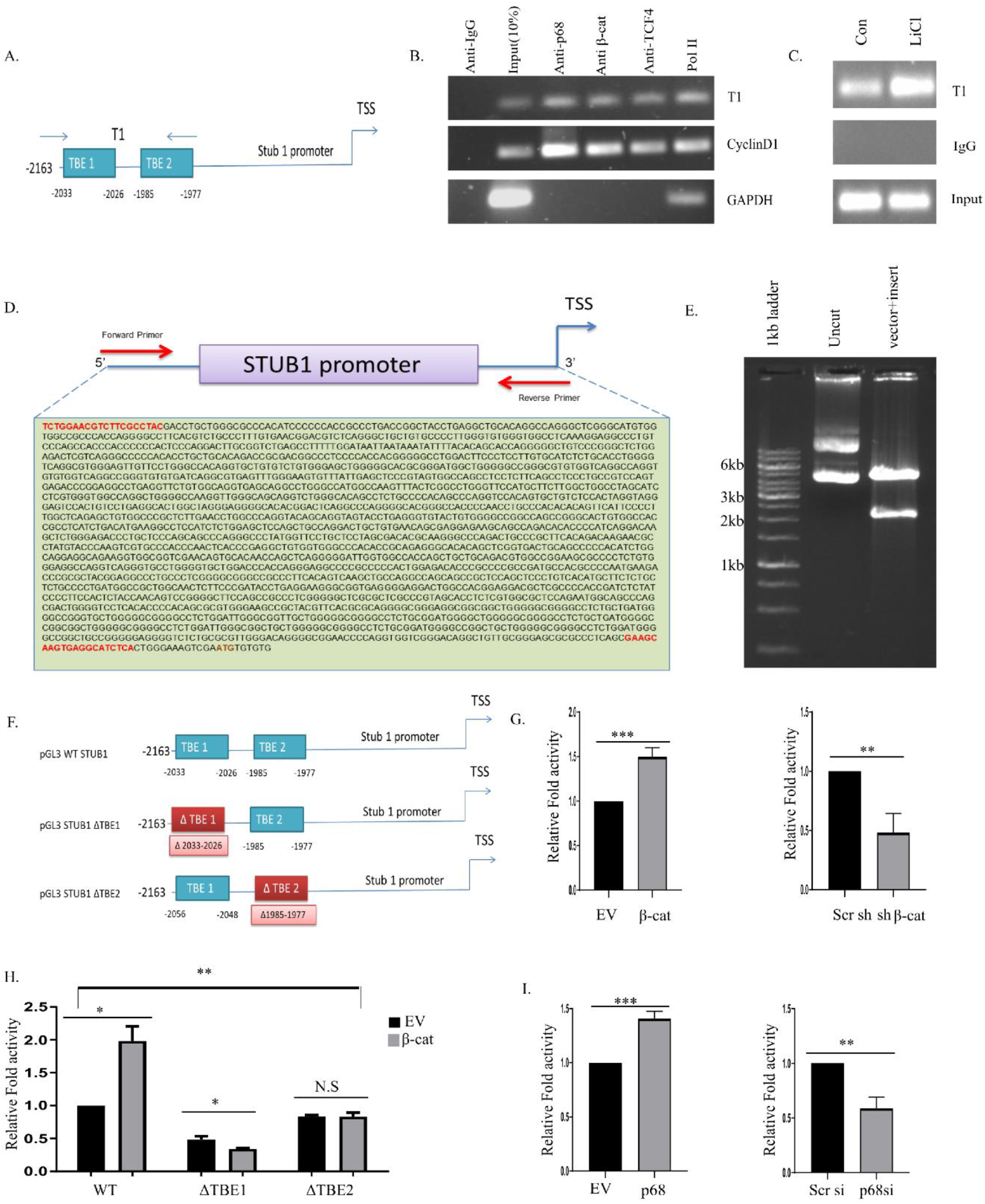
Transcriptional activation of CHIP (*STUB1*) gene promoter is under the control of both β-catenin and p68. *A,* Schematic representation of *STUB1* promoter indicating TCF-binding Elements (TBE1 and TBE2) and Transcription Start Site (TSS). *B,* HCT 116 cells were subjected to Chromatin Immunoprecipitation (ChIP) assay using the denoted antibodies. Amplification of promoter region of Cyclin D1 containing TBE acted as positive control for both β-catenin as well as p68. GAPDH promoter was used as a positive control for RNA Polymerase II. Immunoprecipitated DNAs were PCR amplified using primers designed from the *STUB1* promoter region containing TBEs (T1). ChIP using RNA pol II and IgG antibodies were used as positive and negative controls, respectively. DNA extract (10% without ChIP) was used as input. *C,* ChIP assay was performed from HCT 116 cells following 30mM LiCl treatment (4 hours). Immunoprecipitated DNAs were PCR amplified using primers designed from the *STUB1* promoter region containing TBEs (T1) as shown in the schematic diagram. ChIP using antibody against IgG served as negative control. DNA extract (10% without ChIP) was used as input. *D,* Nucleotide sequence of *STUB1* promoter with the TBEs and TSS highlighted. *E, STUB1* promoter was cloned in pGL3 basic vector. *F,* Schematic representation of WT *STUB1* promoter and its deletion mutants, ΔTBE1 (Δ2033-2026) and ΔTBE2 (Δ1985-1977) sub-cloned in pGL3 basic vector. *G,* HCT 116 cells were transfected with either EV or β-catenin (left) and either Scrambled shRNA or β-catenin shRNA (right), along with pGL3-WT-*STUB1* promoter and Renilla luciferase plasmid (50ng). *H,* HCT 116 cells were transfected with either pGL3-WT-*STUB1* promoter, or its deletion mutants, pGL3-*STUB1*-ΔTBE1 and pGL3-*STUB1*-ΔTBE2, along with Renilla luciferase plasmid (50ng) and EV or β-catenin. *I,* HCT 116 cells were transfected with either EV or p68 (left) and either Scrambled siRNA or p68 siRNA (right), along with pGL3-WT-*STUB1* promoter and Renilla luciferase plasmid (50ng). For *G, H,* and *I,* Renilla luciferase activity was designated for normalization and the acquired data represented as fold activity with respect to control. *Con:* Control, *β-cat:* β-catenin *Scr sh:* Scrambled shRNA, *shβ-cat:* β-catenin shRNA, *Scr si:* Scrambled siRNA. Error bars in all the designated subfigures represent mean (+) s.d. from independent biological repeats. Indicated p values were calculated using Student’s t-test and p≤0.00005 is represented as ****, *NS:* Not Significant.

### β-catenin-mediated enhanced expression of CHIP augments cellular proliferation and survival in colorectal cancer cells thus boosting oncogenesis

The Wnt/β-catenin signaling pathway is known to be abnormally upregulated in colorectal cancer and thus aids in its cellular proliferation and poor prognosis of the cancer.^44^ Also, from the above findings, it has been established that CHIP expression is upregulated by this pathway. To check the effects of the ‘β-catenin-CHIP signaling’ axis on cellular proliferation, clonogenic assays were done after enhancing the effect of β-catenin by LiCl treatment. An increase in the number of colonies was observed with LiCl treatment as compared to the control (Figure 6A). Likewise, when CHIP levels were enhanced in the cell by overexpression of the protein directly in the cells, a similar change was observed (Figure 6A). The culture plate comprising of CHIP overexpression, had a much larger number of colonies compared to the control. These were converted to percentage change in the number of colonies and represented as bar graphs (Figure 6A). When β-catenin signaling was inhibited by FH535, a significant reduction in the number of colonies was obtained (Figure 6B). Similar results were observed when CHIP was knocked down *via* transfection of its siRNA (Figure 6B). A substantial decrease in the number of colonies obtained in both cases was represented as bar diagram (Figure 6B). Thus, the number of colonies increased when both β-catenin pathway and CHIP were enhanced and decreased when they were reduced. MTT assays also revealed that both LiCl-mediated nuclear β-catenin stabilization and CHIP overexpression led to an increase in the viability of HCT 116 cells, while upon diminishing the Wnt/β-catenin pathway by FH535 or upon knocking down CHIP, the percentage viability of HCT 116 cells decreased (Figure 6C). Further, wound healing assay was performed after LiCl treatment, which showed an increased percentage of healing of the wound after 24 hours of the treatment thus highlighting that the treatment aids in cellular proliferation (Figure 6D). Likewise, cells transfected with CHIP also revealed faster healing than its control when observed after 24 hours of transfection (Figure 6E). A significant decrease in the wound was represented in the graphs 24 hours post LiCl treatment or CHIP transfection compared to their respective controls (Figure 6D, 6E). Wound healing assays were also performed after FH535 treatment (Figure S2A) which yielded decreased healing of the wound after 24 hours as observed from a lesser filling of the gap in the FH535 treated cells compared to the control. This was also demonstrated in the bar graphs (Figure S2A). Additionally, when CHIP was knocked down by siRNA, there was a similar observation of less wound healing compared to its control, represented by its respective bar graph (Figure S2B).

**FIGURE 6.**
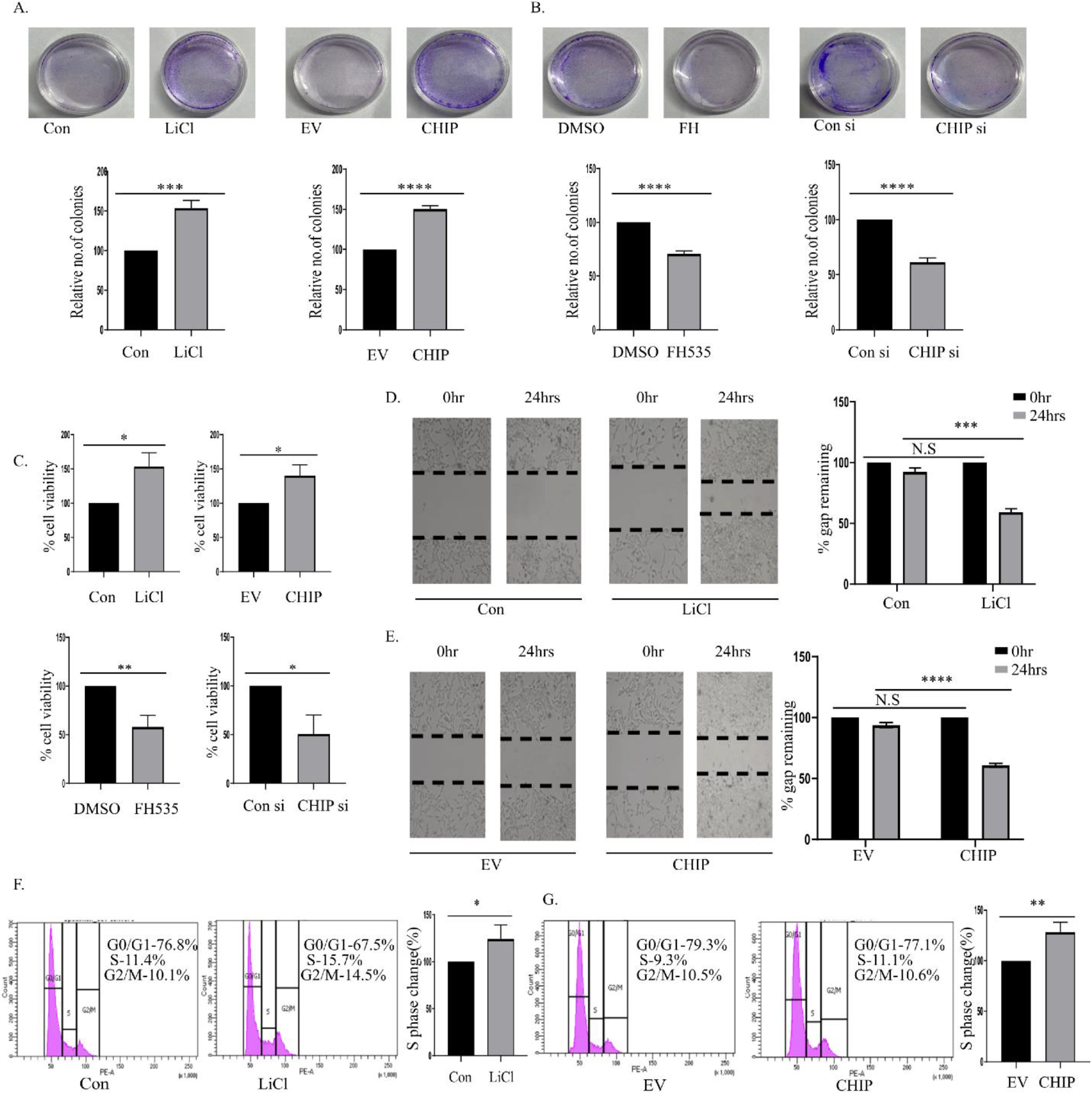
β-catenin-mediated enhanced expression of CHIP augments cellular proliferation and survival in colorectal cancer cells thus boosting oncogenesis. *A,* HCT 116 cells treated with control or 30mM LiCl (4 hours) (left), or alternatively, transfected with 2ug of empty vector or CHIP (48 hours) (right) were checked for their colony formation ability. After 2 weeks, the plates with colonies were stained with crystal violet, kept overnight at 4°C before counting. Bar diagram represents the relative number of colonies averaged from three biological repeats. *B,* HCT 116 cells treated with DMSO or 15uM FH535 (24 hours) (left), or alternatively, transfected with 2ug of Scrambled siRNA or CHIP siRNA (48 hours) (right) were checked their ability to form colonies. After 2 weeks, the colonies were stained with crystal violet, kept overnight at 4°C before counting. Bar diagram represents the relative number of colonies averaged from three biological repeats. *C,* HCT 116 cells treated with control or 30mM LiCl (4 hours) (upper left), or alternatively, transfected with 2ug of EV or CHIP (48 hours) (upper right) were checked for cell viability using MTT assay. HCT 116 cells treated with DMSO or 15uM FH535 (24 hours) (lower left), or alternatively, transfected with 2ug of Scrambled siRNA or CHIP siRNA (48 hours) (lower right) were checked for their viability through MTT assay. The average absorbance values of three biological repeats were plotted. *D,* HCT 116 cells seeded in 35 mm cell culture dishes were treated with control or 30mM LiCl (4 hours). After making transverse scratches on the plates with 10ul sterile pipette tips, the images were captured at the specified time points. Scale bar-400um. Bar diagram represents percentage of gap remaining. *E,* HCT 116 cells seeded in 35 mm cell culture dishes were transfected with 2ug of EV or CHIP. After making transverse scratches on the plates with 10ul sterile pipette tips, the images were captured at the specified time points. Scale bar-400um. Bar diagram represents percentage of gap remaining. *F,* Cell cycle distribution pattern of HCT 116 cells treated with control or 30mM LiCl (4 hours) was checked by flow cytometry. Bar graph represents the change in percentage of S phase cells in the treated compared to the control. *G,* Cell cycle distribution pattern of HCT 116 cells transfected with EV or CHIP was checked by flow cytometry. Bar graph highlights the change in percentage of S phase cells. *Con:* Control, *Scr si:* Scrambled siRNA, *EV:* Empty Vector, *NS:* Not Significant. Error bars in all the indicated subfigures represent mean (+) s.d. from three independent biological repeats. Indicated p values were calculated using Student’s t-test and p≤0.00005 is represented as ****.

The β-catenin-CHIP network also led to an alteration in the cell cycle profile. An increase in the percentage of S phase cells was observed upon LiCl treatment (Figure 6F) or CHIP overexpression (Figure 6G). In contrast, a decrease in the percentage of S phase cells was observed on treating cells with FH535 (Figure S2C). Collectively, the enhancement of CHIP *via* the Wnt/β-catenin signaling pathway fortifies cellular proliferation and enhanced migration in colorectal cancer cells.

## DISCUSSION

Cancer is one of the chief reasons for ill health and mortality in the world ^45^, necessitating a complete understanding of the detailed mechanism of oncogenesis. Despite various approaches to elucidate different aspects of disease progression, the understanding remains incomplete.

CHIP is an E3 ubiquitin ligase known to maintain protein homeostasis in cells by degrading several proteins in different disease contexts. Although upregulated in colon cancer, prostate cancer, and lung cancer ^12,16,17^, CHIP shows much lower expression in breast cancer and glioma ^18,19^. It is known to target both tumor suppressors and oncogenes ^46^ in different cancers but the exact mechanism of this is unclear. Thus, CHIP expression and its degradation of substrate proteins are largely context-dependent. The reasons underlying this differential expression of CHIP remain poorly understood. Keeping the existing literature in view, we sought to understand how the expression of CHIP is regulated, and in this context, colorectal cancer was chosen. From a recent report stating how CHIP functions as an oncogene in colorectal cancer ^12^ and the Human Protein Atlas, CHIP was found to have a higher level of expression in colorectal cancer. Bioinformatic analyses from available web-based tools like UALCAN ^30^. HPA ^13^ and Oncomine ^32^ revealed an elevated expression of CHIP (*STUB1*) gene in colorectal cancer as indicated in Figure1. This led to further explorations into the mechanistic detail of this increased level of CHIP expression, which was corroborated with detailed experimental results. The major regulator of colorectal cancer prognosis is the Wnt/β-catenin signaling pathway which, concomitant with elevated transcription of cancer-promoting genes, leads to a poor prognosis. Here, we focused on how this pathway might aid in the transcription of CHIP (*STUB1)* gene. Initial treatments with the ligand Wnt3a, which triggered the Wnt/β-catenin signaling pathway, exhibited an increase in CHIP mRNA and protein levels in treated colorectal cancer cell HCT 116 compared to its respective control. Following this, colorectal cancer cell lines were treated with activators or inhibitors of the Wnt/β-catenin signaling pathway. Upon activating the pathway, an increase in CHIP protein level was observed while on inhibiting the pathway, CHIP protein level diminished. Immunoblot analysis upon LiCl treatment to enhance nuclear translocation of β-catenin revealed an increased amount of CHIP protein in both the cytoplasmic and nuclear fractions. As CHIP is majorly a cytoplasmic protein, an increased nuclear accumulation requires further study into the matter, especially its subsequent function in the nuclear compartment in the context of colorectal cancer. To strengthen our hypothesis that CHIP expression is dependent on Wnt/β-catenin, β-catenin was overexpressed or knocked down in colorectal cancer cell lines and CHIP expression was observed at both mRNA and protein levels. Upon overexpressing β-catenin, CHIP mRNA and protein levels were elevated and on knocking down β-catenin, they were reduced. A positive correlation was found, suggesting that CHIP mRNA was positively regulated by β-catenin, and this effect was transduced to the protein level.

p68 is often involved in transcriptional activation of β-catenin-driven genes in several cancers including colorectal cancer ^8,42^. To investigate any possible role of p68 in CHIP gene transcription, p68 levels were modulated. p68, being a co-activator, also positively correlated with CHIP mRNA and protein levels. Upon simultaneous elevation of p68 and β-catenin pathway components, a higher expression level of CHIP protein was observed compared to their individual overexpression. Also, when rescue experiments were performed by knocking down β-catenin and subsequently upregulating p68, CHIP protein levels, that had decreased on the former condition, were somewhat restored in the rescue condition *i.e.,* upon p68 overexpression. Thus, our results establish that p68 work in conjunction with the β-catenin pathway for regulation of CHIP gene expression.

To delve a little deeper into the transcriptional control of *STUB1* mRNA, we cloned the CHIP (*STUB1*) promoter. Bioinformatics studies conducted on the available promoter sequence revealed the presence of putative TCF4 binding sites, through which β-catenin binds and regulates transcription of its target genes. A 2163 bps sequence of the promoter containing the TCF4 binding sites was cloned. This is the very first report of *STUB1* promoter cloning, which was further used to study the promoter strength upon induction of transcription factors that enhance *STUB1* transcription. ChIP studies revealed the binding of β-catenin, p68, and TCF4 on the CHIP promoter. The promoter strength, measured by fold change of activity *via* luciferase assay, ascertained the association of p68 and β-catenin in positively regulating CHIP promoter activity. This was further corroborated with studies where the TCF4 binding sites on the *STUB1* promoter were mutated, and a reduction of promoter activity was observed compared to the wild type upon β-catenin overexpression. Various assays established the physiological implications of this increase in CHIP mRNA and protein levels, divulging the fact that CHIP aids in increased cellular proliferation and migration, which is in accordance with the role of the β-catenin pathway in colorectal cancer. A schematic representation of the Wnt/β-catenin-p68-mediated mechanism of CHIP gene regulation and its outcome towards uncontrolled cellular propagation in colorectal cancer has been portrayed in Figure 7.

**FIGURE 7.**
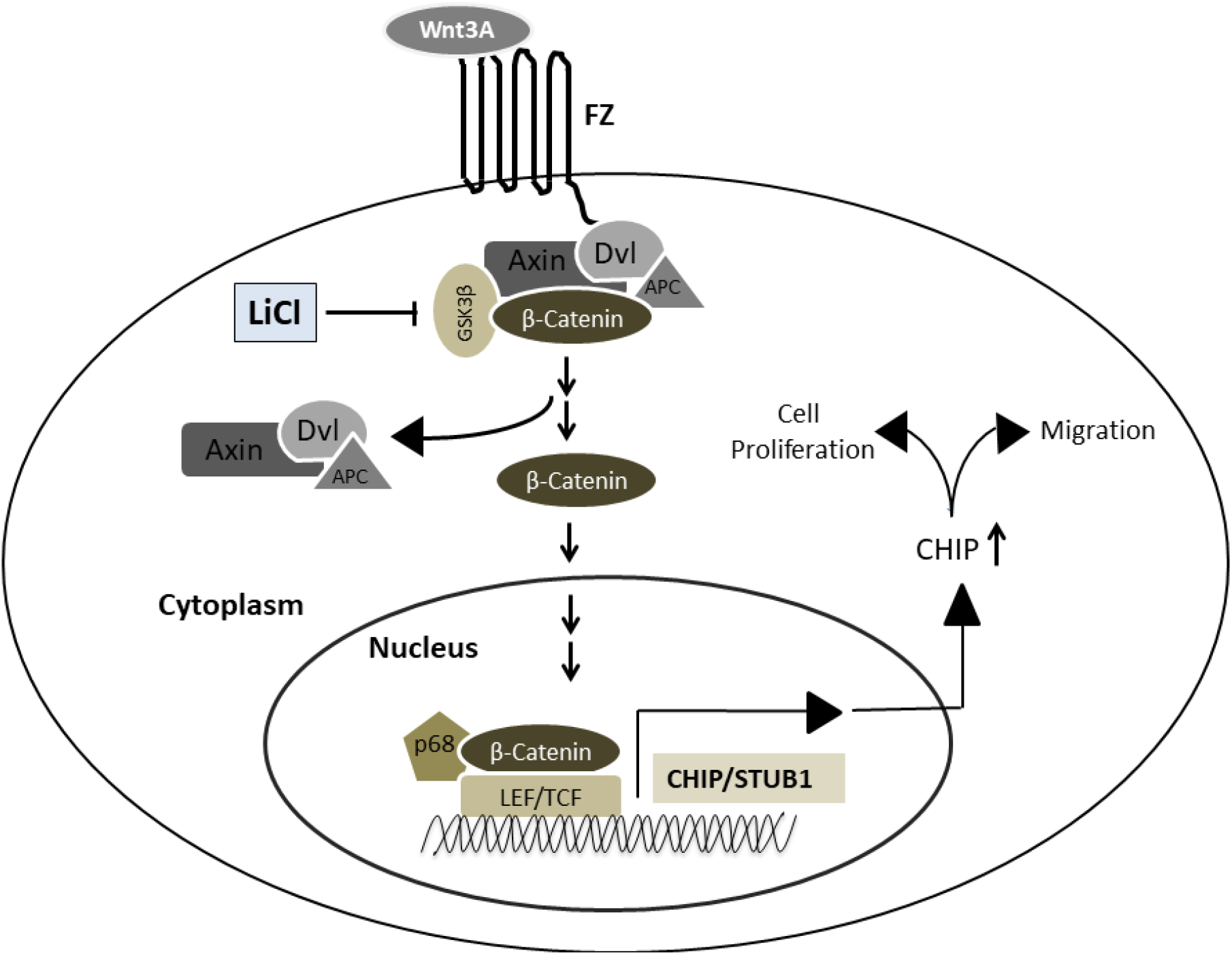
Schematic representation of Wnt/β-catenin and p68 mediated transcription of CHIP gene. From the current study a model is proposed which states that upon stabilization by Wnt signaling agonists like LiCl, Wnt3a and upon overexpression, β-catenin occupies the CHIP promoter and leads to the latter’s increased transcription. This is augmented by RNA helicase p68, the co-activator of β-catenin. The resultant increased CHIP gene expression leads to enhanced cellular proliferation, cell survival, and migration of colorectal cancer cells.

In a contrasting report of CHIP acting as a tumor suppressor in colorectal cancer, decreased level of CHIP protein was observed in 37.2% of the patient samples ^47^. However, 62.8% of the samples revealed an increased or unchanged CHIP protein level, demanding further study.

Thus, in the current study, we elucidated the hitherto unclear regulation of CHIP gene expression by establishing a Wnt signaling pathway-driven mechanism of CHIP expression in colorectal cancer. Although different pathways need to be explored in different contexts, this initial report will provide a basis for future studies on the gene-level regulation of this pivotal E3 ligase that can affect multiple cancers. Additionally, this study highlights the necessity of monitoring CHIP expression levels while making therapeutic approaches to curb colorectal cancer.

## Supporting information

BioRxiv_Supplementary file_Ghosh MK

