## Supplementary material for "Wnt/β-catenin signaling and p68 conjointly regulate CHIP in colorectal cancer": BioRxiv_Supplementary file_Ghosh MK: Supplementary files - a single pdf_Ghosh MK.pdf

**Appendix A: List of primers used.**

| Name | Sequence (5'-3') |
|---|---|
| **Cloning Primer** | |
| *STUB1* (CHIP) promoter | F-5'-AATAGATCTTCTGGAACGTCTTCGCCTAC-3'<br>R-5'-AATAAGCTTTGAGATGCCTCACTTGCTTC-3' |
| **qRT-PCR Primers** | |
| *STUB1* (CHIP) | F-5′-AGGCCAAGCACGACAAGTACAT-3′<br>R-5′-CTGATCTTGCCACACAGGTAGT-3' |
| *CCND1* (Cyclin D1) | F-5'-CCGTCCATGCGGAAGATC-3'<br>R-5'-GAAGACCTCCTCCTCGCACT-3' |
| *DDX5* (p68) | F-5'-TGAGCGACCTTATCTCTGTGC-3'<br>R-5'-CCTGGAACGACCTGAACCTC-3' |
| *CTNNB1* (β-catenin) | F-5'-TACCTCCCAAGTCCTGTATGAG-3'<br>R-5'-TGAGCAGCATCAAACTGTGTAG-3' |
| *18S rRNA* | F-5'-GCTTAATTTGACTCAACACGGGC-3'<br>R-5'-AGCTATCAATCTGTCAATCCTGTC-3' |
| **ChIP Primers** | |
| T1-TCF4 binding element on *STUB1* promoter | F -5'-CAGGCCAGGGCTCGGGC -3'<br>R-5' -CCGCAAGTCCTGGGAGTG -3' |
| *Cyclin D1-ChIP* | F-5'-GTAACGTCACACGGACTACAGG-3'<br>R-5'-GCACACATTTGAAGTAGGACACC-3' |
| *GAPDH-ChIP* | F-5'- CGGCTACTAGCGGTTTTACG-3'<br>R-5'- GGCTGCGGGCTCAATTTAT-3' |

| Site Directed Mutagenesis Primers | |
|---|---|
| ΔTBE1 | F- 5’-CCTTCACGTCTGCCACGGACGTCTCAGG-3’<br>R-5’-CCTGAGACGTCCGTGGCAGACGTGAAGG-3’ |
| ΔTBE2 | F - 5’-GGGTGTGGGTGGGAGGCCCTGTCC-3’<br>R- 5’-GGACAGGGCCTCCCACCCACACCC-3’ |

### Supplementary figures and figure legends

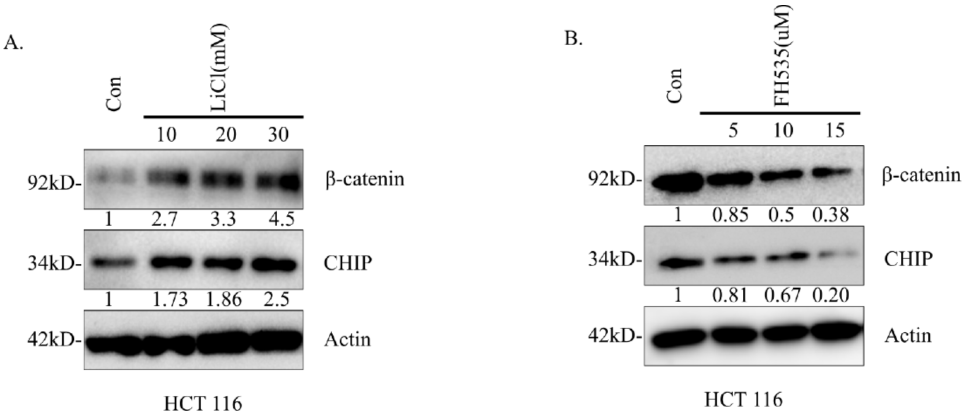


**FIGURE S1.** *A,* HCT 116 cells were administered with increasing concentration of LiCl (10mM, 20mM and 30mM respectively) in a dose dependent manner for 4 hours along with control. The prepared whole cell lysates were immunoblotted and probed for proteins β-catenin and CHIP with their respective antibodies. *B,* HCT 116 cells treated with increasing concentration of FH535 (5uM, 10uM and 15uM respectively) in a dose dependent manner along with DMSO control for 24 hours. *Con:* Control. Densitometric values were assigned with respect to loading control Actin.

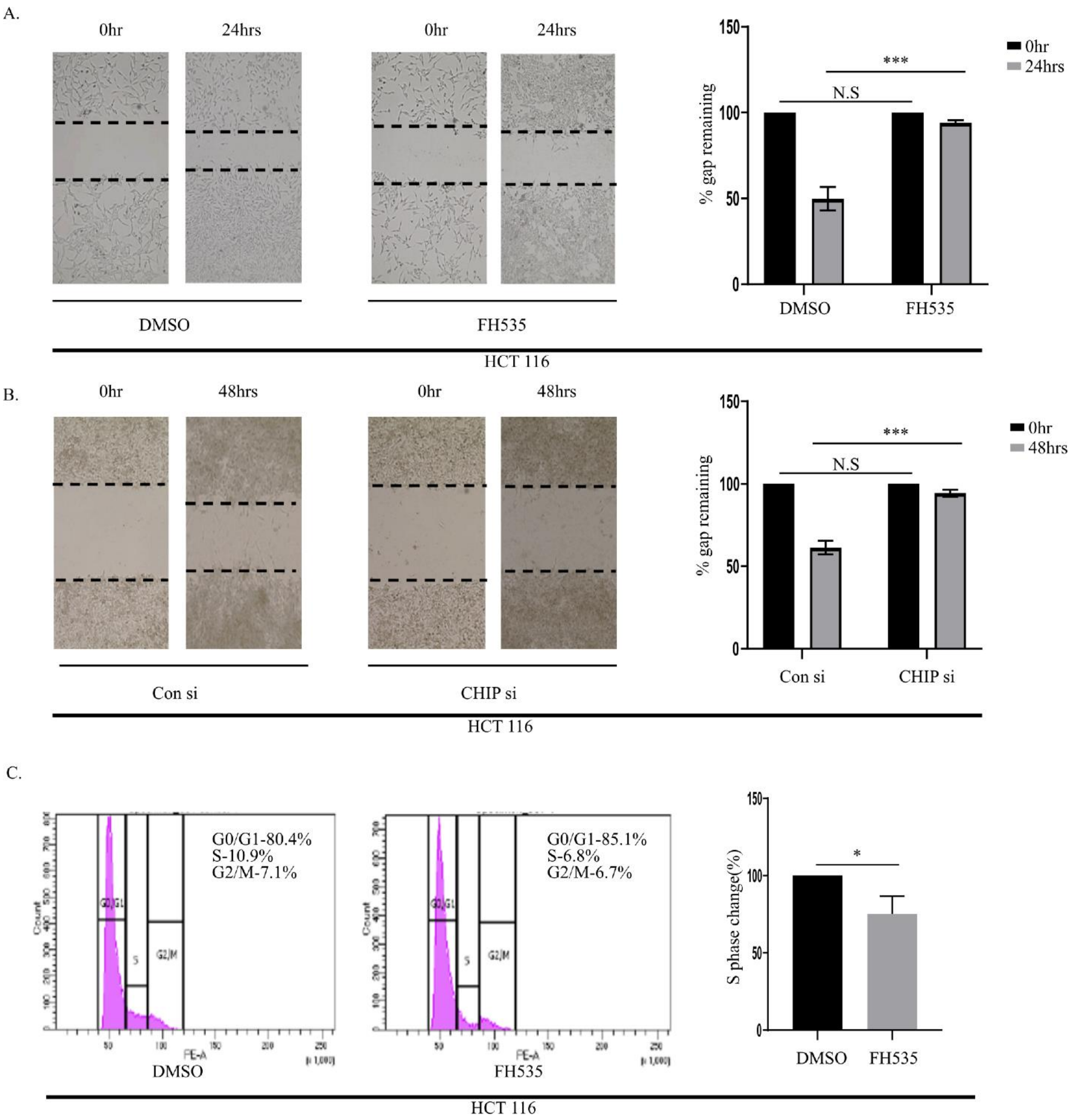


**FIGURE S2.** *A*, HCT 116 cells seeded in 35 mm cell culture dishes were treated either with DMSO control or 15uM FH535 (24 hours). After making transverse scratches on the plates using 10ul sterile pipette tips, the cells were grown for another 24 hours before the images were captured at the time points specified in the figure. Scale bar-400um. Bar diagrams represent the percentage of gap remaining. *B*, HCT 116 cells seeded in 35 mm cell culture dishes were transfected with Scrambled siRNA or CHIP siRNA. After making transverse scratches on the plates with 10ul sterile pipette tips, the cells were grown for another 24 hours before the images were captured at the time points specified in the figure. Scale bar-400um. Bar diagrams represent the percentage of gap remaining. *C*, HCT 116 cells treated with DMSO control or 15uM FH535 (24 hours) were subjected to flow cytometry and cell cycle distribution pattern was checked. Bar graphs represent the change in the percentage of S phase cells in the treated compared to the control. Error bars in all the designated subfigures represent mean (+) s.d. from three independent biological repeats. Indicated p values were calculated using Student's t-test and $p \leq 0.00005$ is represented as ****.
